# Cryo-EM structure of alpha-synuclein fibrils

**DOI:** 10.1101/276436

**Authors:** Ricardo Guerrero-Ferreira, Nicholas M. I. Taylor, Daniel Mona, Philippe Ringler, Matthias E. Lauer, Roland Riek, Markus Britschgi, Henning Stahlberg

**Author notes:** Correspondence to: Henning Stahlberg. Current Address: Structural Biology of Molecular Machines Group, Protein Structure & Function Programme, Novo Nordisk Foundation Center for Protein Research, Faculty of Health and Medical Sciences, University of Copenhagen, Blegdamsvej 3B, Copenhagen 2200, Denmark (Nicholas M. I. Taylor).

## Abstract

Intracellular inclusions of alpha-synuclein are the neuropathological hallmark of progressive disorders called synucleinopathies. Alpha-synuclein fibrils are associated with transmissive cell-to-cell propagation of pathology. We report the structure of an alpha-synuclein fibril (residues 1-121) determined by cryo-electron microscopy at 3.4Å resolution. Two protofilaments form a polar fibril composed of staggered β-strands. The backbone of residues 38 to 95, including the fibril core and the non-amyloid component region, are well resolved in the EM map. Residues 50-57, containing three mutation sites associated with familial synucleinopathies, form the interface between the two protofilaments and contribute to fibril stability. A hydrophobic cleft may have implications for fibril elongation, and inform the rational design of molecules for diagnosis and treatment of synucleinopathies.

## Introduction

Parkinson’s disease (PD) is a neurodegenerative disorder characterized by the presence of Lewy bodies (LB) and Lewy neurites (LN), which contain the presynaptic protein alpha-synuclein (α-Syn, 140 residues, ∼14 kD) (Spillantini et al. 1997). Several factors point to α-Syn as an important player in the onset of PD: (i) six known point mutations in the α-Syn gene (SNCA) are associated with familial forms of synucleinopathies (A30P (Kruger et al. 1998); A53T (Polymeropoulos et al. 1997); E46K (Zarranz et al. 2004); H50Q (Appel-Cresswell et al. 2013); G51D (Lesage et al. 2013); A53E (Pasanen et al. 2014)); (ii) animal models suggest a role of α-Syn in the etiology of PD, Dementia with Lewy Bodies (DLB) and Multiple System Atrophy (MSA) (Feany and Bender 2000; Hashimoto, Rockenstein, and Masliah 2003; Periquet et al. 2007; Tyson et al. 2017); (iii) individuals with duplications or triplications of the α-Syn gene exhibit overexpression of α-Syn and develop PD (Singleton et al. 2003; Ibanez et al. 2004).

Certain fibril forms of α-Syn can seed α-Syn inclusions in cell culture and intra-neuronal aggregation of mouse α-Syn in vivo (Luk et al. 2009; Volpicelli-Daley, Luk, and Lee 2014), which makes α-Syn fibrils an important target for the development of diagnostic tools and therapeutic strategies for PD and related synucleinopathies. However, high resolution structures of α-Syn fibrils are limited to the results of a micro-electron diffraction (microED) study of small segments of the protein (Rodriguez et al. 2015) and a solid-state NMR structure of ∼5 nm diameter, single protofilaments (Tuttle et al. 2016), in addition to solid state NMR studies at the secondary structure level (Vilar et al. 2008; Bousset et al. 2013; Kim et al. 2009).

Because understanding the structure of α-Syn fibrils at high resolution is pivotal to realize their mechanism of nucleation and propagation, we report here the atomic structure of human α-Syn fibrils determined by cryo-electron microscopy (cryo-EM). Our results could inform strategies to inhibit fibril formation and the development of small molecules for the diagnosis and treatment of synucleinopathies. We provide the structural basis of the organization of α-Syn fibrils at near-atomic resolution and suggest a mechanism of fibril formation, stability and growth.

## Results and Discussion

### The 3D structure of α-Syn amyloid fibrils

Several preparations of recombinant human α-Syn fibril were screened by negative stain transmission electron microscopy (TEM; Figure 1 – figure supplement 1). These included fibrils formed by full length α-Syn (Figure 1A), α-Syn phosphorylated at serine 129, and by C-terminal truncated α-Syn comprised of residues 1-119 (α-Syn(1-119)), 1-121 (α-Syn(1-121)), or 1-122 (α-Syn(1-121)). The fibrils formed by α-Syn(1-121) were straight, between 20 and 500 nm long and the only ones of consistently 10 nm in diameter (Figure 1 – figure supplement 1E). For this reason, α-Syn(1-121) was used to proceed with structural analysis by cryo-EM (Figure 1B). Helical image processing produced a 3D reconstruction of the fibril at an overall resolution of 3.4 Å (Figure 1C and D, Figure 1 – figure supplement 2, Figure 2 and Video 1).

**Figure 1.**
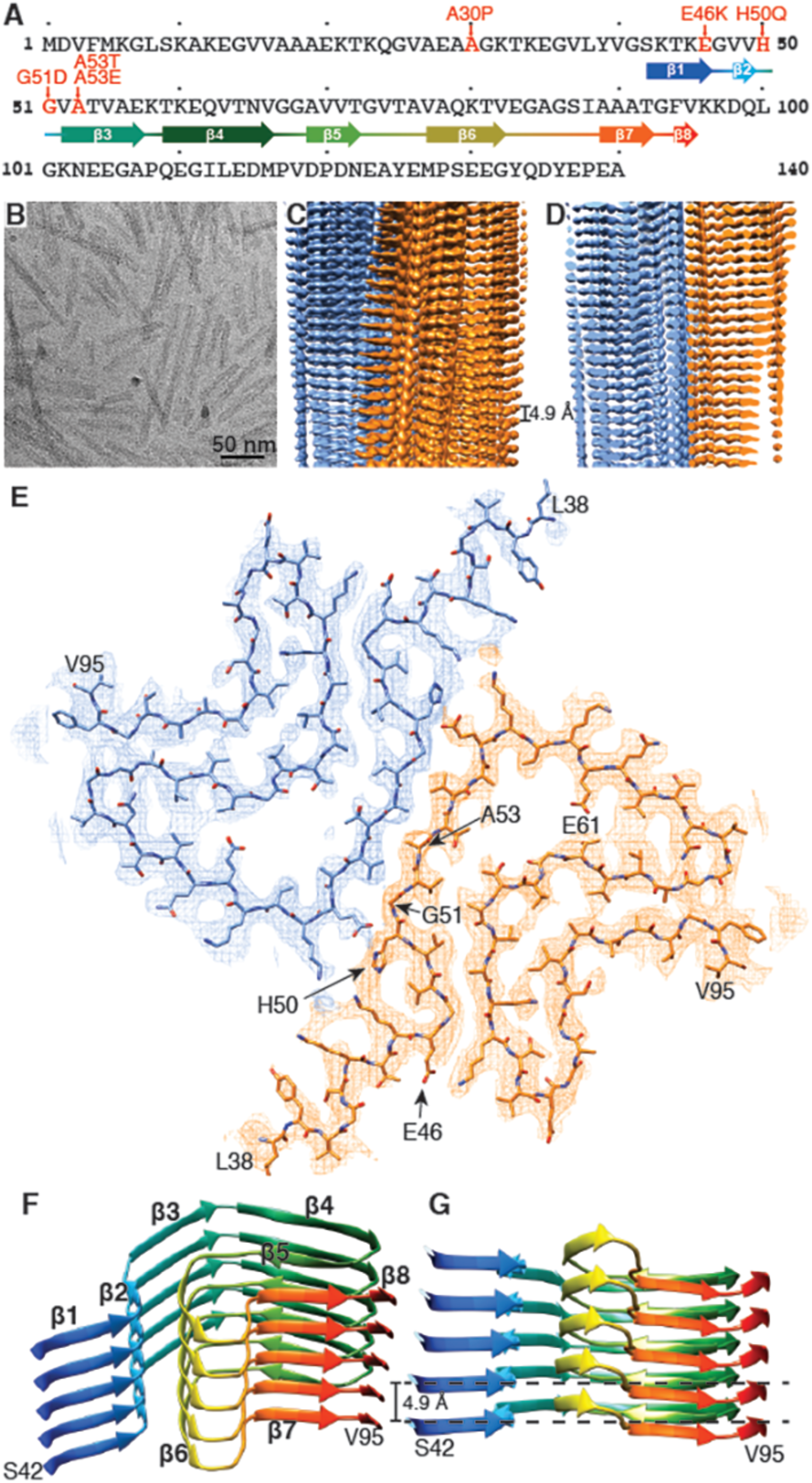
Structure of α-Syn(1-121) fibril. **(A)** Schematic depicting the sequence of human α-Syn. The positions of the known familial mutations are indicated. β-strand regions are indicated by arrows colored from blue to red. **(B)** Cryo-EM micrograph depicting the distribution and general appearance of α-Syn fibrils. **(C)** Cryo-EM reconstruction of α-Syn(1-121) fibrils showing two protofilaments (orange and blue). **(D)** Cross-section of (C) illustrating the clear separation of the β-strands, also shown in Figure 1 – figure supplement 3A and B. **(E)** Cross-section of a fibril (along the axis) illustrating the arrangement of the two protofilaments (orange and blue) and fitted atomic model. Positions of the initial (L38) and final (V95) residues fitted are indicated, as well as the initial and final residue of the NAC region (E61 to V95). Arrows indicate the location of four of the five α-Syn residues where familial mutations associated with PD occur. **(F)** Distribution of β-strands in a single protofilament of the α-Syn fibril, corresponding to residues 42 to 95. Color scheme, as in (A). **(G)** As in (F) but a perpendicular view to the fibril axis illustrating height differences in some areas of a single protofilament.

**Figure 2.**
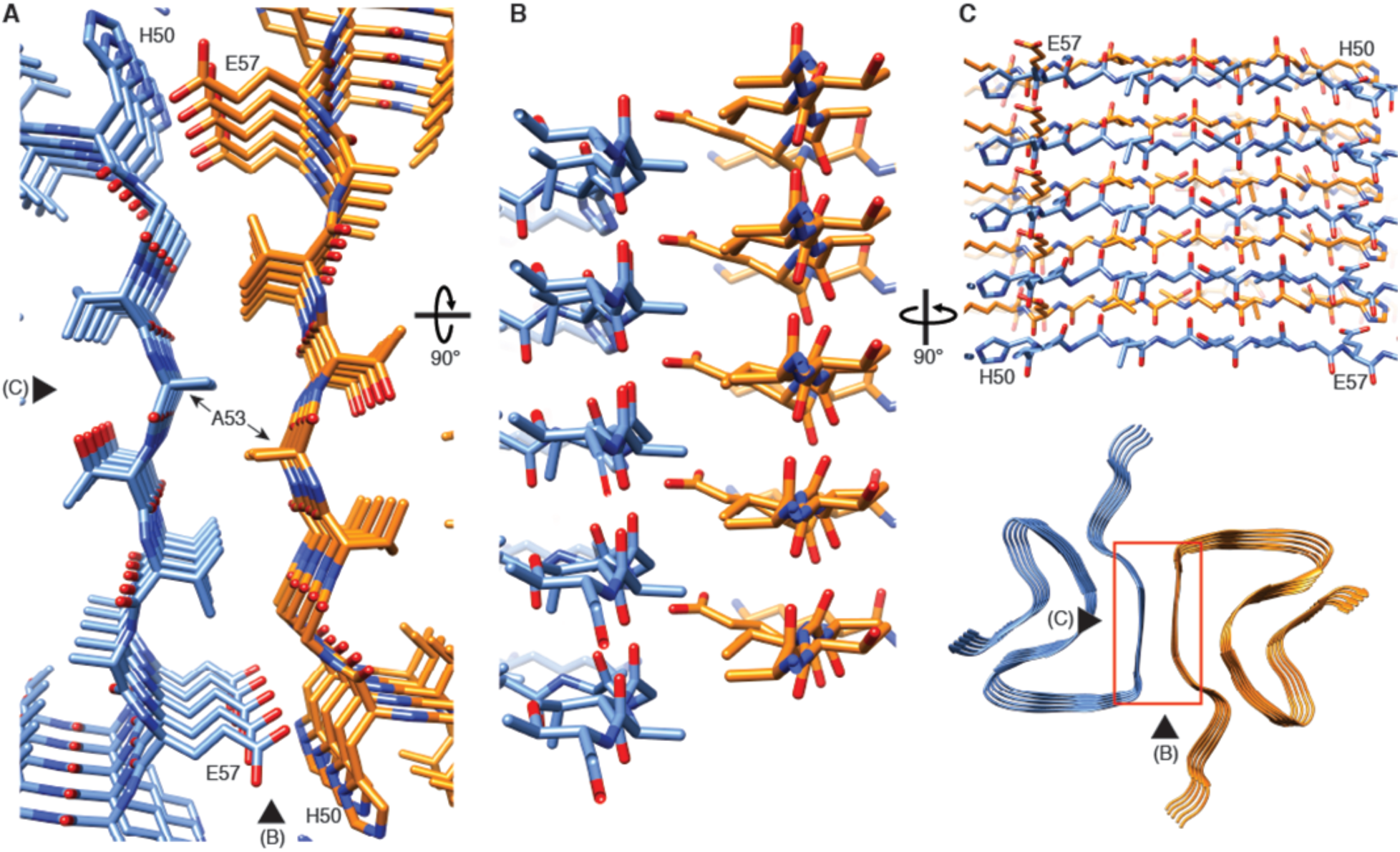
Interface region between two protofilaments of the α-Syn(1-121) fibril. **(A)** View along the axis of the fibril as indicated by the red rectangle on the ribbon diagram (bottom right). **(B) (C)** Side views of the fibril with orientations indicated by arrowheads in (A) and the ribbon diagram (bottom right). Panels (B) and (C) clearly illustrate the 2_1_ screw symmetry that results from the staggered arrangement of subunits.

The 3D map shows that α-Syn(1-121) fibrils are formed by two protofilaments of 5 nm in diameter (Figure 1). These lack C2 symmetry, but are related by a 2_1_ screw symmetry akin to the screw symmetry exhibited by the paired helical filaments of tau (Fitzpatrick et al. 2017) and by amyloid-ℬ(1-42) filaments (Gremer et al. 2017). α-Syn(1-121) fibrils are therefore polar. The position of a given ℬ-sheet in a protofilament is produced by the rotation of −179.5° of one sheet around its axis (helical twist), followed by a vertical translation of 2.45 Å (helical rise). This ℬ-sheet arrangement results in a spacing of 4.9 Å between α-Syn subunits in successive rungs of a single protofilament (Figure 1C and D). The quality of the EM map allowed an atomic model of the region between residues L38 and V95 to be built.

Each α-Syn(1-121) molecule comprises eight in-register parallel β-strands (*i.e.*, residues 42-46 (β1), 48-49 (β2), 52–57 (β3), 59–66 (β4), 69-72 (β5), 77–82 (β6), 89-92 (β7), and 94-(∼102) (β8)), which are interrupted by glycine residues (*i.e.*, G41 before β1, G47 between β1 and β2, G51 between β2 and β3, G67 and G68 between β4 and β5, G73 between β5 and β6, G84 and G86 between β6 and β7, and G93 between β7 and β8) or an arch (*i.e.*, E57-K58 between β3 and β4) (Figure 1A, F, and G). The β-strands β2-β7 wind around a hydrophobic intra-molecular core composed of only alanine and valine residues and one isoleucine (*i.e.*, V48, V49, V52, A53, V55, V63, A69, V70, V71, V74, A76, V77, A78, I88, A89, A90, A91) in an organization resembling a Greek key-like motif as recently suggested (Figure 1A and Figure 3) (Tuttle et al. 2016). Considering that these hydrophobic clusters are maintained along the fibril, they are likely to contribute to the stability of the protofilament. The hydrophobic core is surrounded by two hydrophilic regions (*i.e.* (i): Q79, T81, and (ii): T72, T75, T54, T59, and E61) both still within the core of the structure (Figure 3). While most of these side chains form so-called side chain hydrogen bond ladders (Riek 2017; Nelson et al. 2005), the second hydrophilic region comprising four threonine residues and a negatively charged glutamic acid side chain surrounds a tunnel filled with some ordered molecules of unknown nature, as evidenced by an electron density (Figure 1-figure supplement 3D). The less well defined β1 and β8 strands are attached to the core, while the first 37 N-terminal residues and the last ∼20 C-terminal residues of α-Syn(1-121) are not visible in the electron density (Figure 1E and Figure 1 – figure supplement 2A), indicating an ill-defined structure in line with quenched hydrogen/deuterium exchange – solution-state NMR (H/D exchange NMR) and limited proteolysis (Vilar et al. 2008), which showed these terminal segments to be unprotected in nature. Together with our results, this suggests that approximately 40 residues of both the N-and C-terminals of full-length human α-Syn are flexible, and surround the structured core of the fibril with a dense mesh of disordered random-coil-like tails.

**Figure 3.**
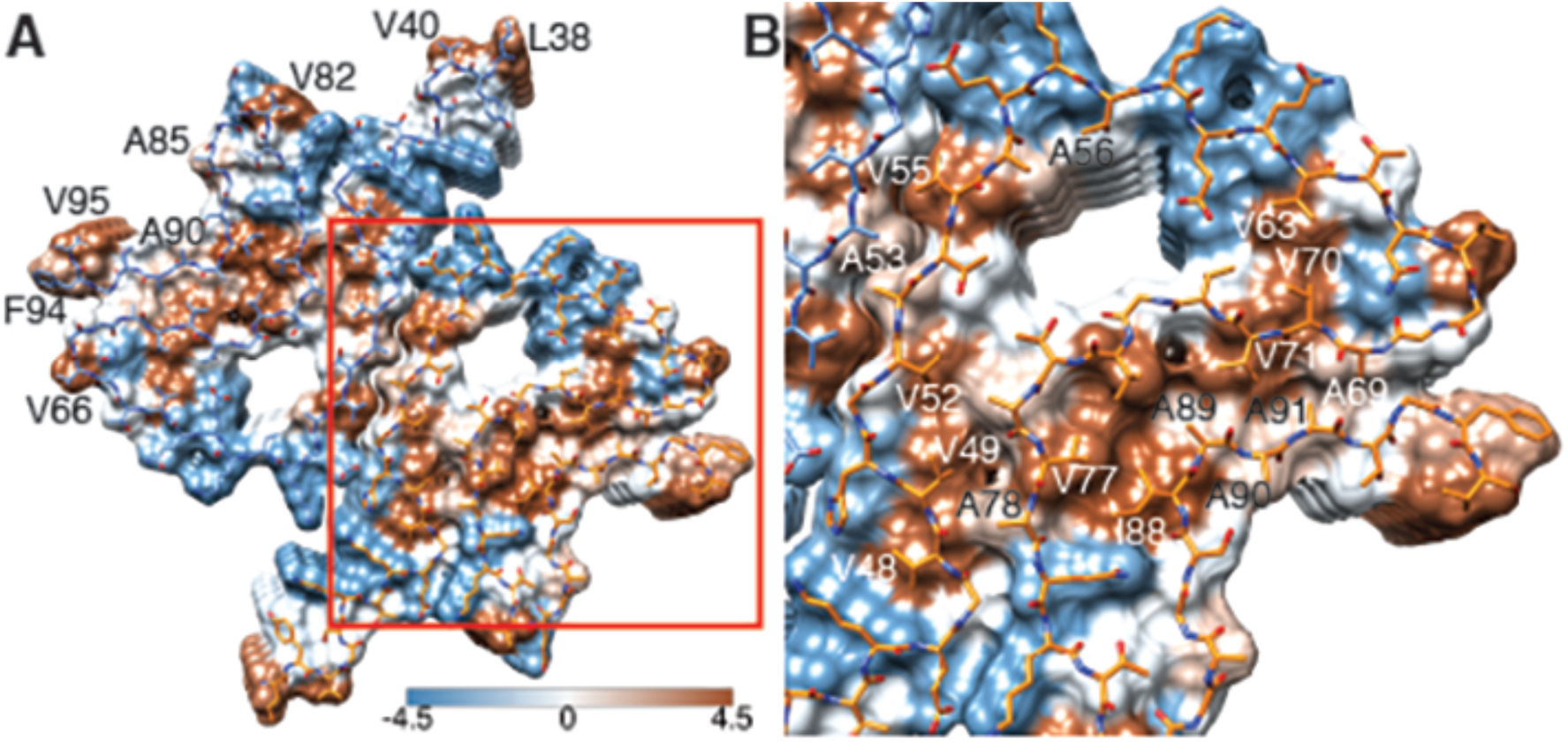
Hydrophobicity of α-Syn(1-121) fibrils. **(A)** Top view (fibril axis) of the hydrophobic regions (brown) in a fibril where the hydrophobic pocket at the interface between two protofilaments is evident. Hydrophobicity score from hydrophilic (−4.5, blue) to hydrophobic (4.5, brown) is indicated by the color bar. Hydrophobic residues on the outer surface of the fibril are indicated. **(B)** Close-up of the region highlighted in (A) indicating the hydrophobic core composed of valines and a single isoleucine (I88). Residues forming the hydrophilic region (blue) that surrounds the hydrophobic region of the core are also visible.

Two β-sheets (one from each protofilament) interact at the fibril core via a rather large hydrophobic steric zipper-geometry comprised of β-strand β3 (*i.e.*, residues G51-A56). As a consequence, two α-Syn molecules per fibril layer are stacked along the fibril axis (Figure 2B and C). The side chains of residues A53 and V55 form the inter-molecular surface contributing to the interface between the two protofilaments, which is possibly further stabilized by a surface-exposed potential salt bridge between E57 and H50 that might be sensitive to the pH, as an unprotected histidine has a pK of ∼6.2 (Figure 1 – figure supplement 3H). The same structure with a steric zipper topology was found in micro-crystals of the peptide comprising residues G47-A56 (Rodriguez et al. 2015). Interestingly, within the Greek key-like motif, the β-strand β6 sandwiched between β-strands β2/β3 and β7 is also aligned with a neighboring molecule but shifted by one stack as shown in Figure 1G and Figure 2 – figure supplement 1. Thus, hetero and homo steric zippers are both present in the 3D structure. Of these, the homo steric zipper at the inter-molecular interface has an extensive and well-packed β-strand interface, forming a very densely packed fibril. This stacking generates an asymmetric fibril with two distinct ends. Furthermore, the entire hydrophobic core of the fibril is composed of β-strands that interact with each other in an entirely half-stacked zipper topology, contrasting with the hydrophilic core comprised of β-strands β4 and β5, which are non-stacked (Figure 1G and Figure 2 – figure supplement 1). The latter confirms previous results from site-directed spin labeling experiments, which show that the region including residues 62-67 at the beginning of the non-amyloid component (NAC) region, has a pronounced lack of stacking interactions (Chen et al. 2007).

The outer surface of the fibril is mostly hydrophilic, with a few exceptions (*i.e.*, L38, V40, V82, A85, A90, F94, V95) (Figure 3A). The side chain of V66 should probably not be classified as surface exposed because of the potential interaction with β-strand β8 (Figure 1 – figure supplement 2A). If the influence of the non-polar alanine residues is excluded due to the small size of their side chains, the surface of the fibrils has two highly hydrophobic patches formed by residues L38 and V40, and by residues F94 and V95. Other interesting properties of the surface are the potential salt bridge formed by the side chains of E46 and K80 (Figure 1 – figure supplement 3G) and the rather highly positive clustering of K43, K45, K58, H50 that requests the binding of a counter-ion, as is supported by an observed electron density (Figure 1 – figure supplement 3C).

### The familial PD mutations in the context of the 3D fibril structure

Six familial mutations in α-Syn are known to be associated with PD and other synucleinopathies (*i.e.*, A30P, E46K, H50Q, G51D, A53E, and A53T). Of these, all but A30P are located in the heart of the core of the fibril structure presented here (Figure 1A and E). E46 forms a salt bridge with K80 (Figure 1 – figure supplement 3G). The mutation of glutamic acid to positively charged lysine would thus induce a charge repulsion between β-strands β1 and β5, likely destabilizing α-Syn fibril structure (Tuttle et al. 2016). The familial PD/DLB-causing mutation E46K was found to enhance phosphorylation in mice (Mbefo et al. 2015), and its toxic effect was increased by the triple-K mutation (E35K, E46K, E61K) in neuronal cells (Dettmer et al. 2017).

Previous high-resolution structures of α-Syn were based on small peptides or single protofilaments (Rodriguez et al. 2015; Tuttle et al. 2016). Our results make possible to recognize the structural contribution of some familial mutations to fibril stability, revealing that H50, G51 and A53 are all involved in the inter-molecular contact between two β-sheets from adjacent protofilaments at the core of the fibrils. The mutation H50Q would likely interfere with the potential salt bridge E57-H50 (Figure 1 – figure supplement 3H). The negative side chains at position 51 in the mutation G51D and 53 in the mutation A53E would likely disrupt the steric zipper interaction between the two molecules, while A53T would change the dense hydrophobic packing of the zipper to a looser and partly hydrophilic one. Our structure allows the effect of the A53T mutation to be rationalized. A53 is part of a hydrophobic pocket that not only participates in the interaction between protofilaments, but also contributes to the stability of the fibrils as the hydrophobic interactions exist along the fibril axis. Therefore, the familial mutations at the core of α-Syn fibril would compromise the formation of the fibril structure presented here. This suggests that this fibril structure might have a native function, and that a different fibril structure (*i.e.*, fibril strain) could be involved in the toxicity of the disease.

It seems likely that the fibril structure presented here doesn’t have a native or mechanistic function by itself, since one would expect evolution to have optimized the amino acid sequence to avoid non-functional hydrophobic surface patches (Figure 3), the hydrophilic tunnel (Figure 1 – figure supplement 3D), and the rather peculiar positively charged side chain arrangement comprised of residues K43, K45, K58, H50 (Figure 1 – figure supplement 3C) as is the case for the functional HET-s amyloid structure (Wasmer et al. 2008). However, similar rather unique structural characteristics have been previously observed for Tau, where lysine and tyrosine residues play a similar stabilizing role in the interface region of two protofilaments of the straight filaments (SF) and the area in the center of the protofilaments is dominated by hydrophilic residues (Fitzpatrick et al. 2017).

It is plausible that these structural features might arise because folding to form the amyloid fibril structure is dictated by the need to bury the maximum number of hydrophobic side-chains as efficiently as possible, as is the case in the Aβ(1-42) amyloid fibrils (Gremer et al. 2017). The here presented structure therefore hints at the possibility that a different structure or mechanism might be the causative agent for PD, at least for familial PD. This could be a different fibril strain (Peelaerts et al. 2015), an oligomeric α-Syn intermediate (Winner et al. 2011; Outeiro et al. 2008; Danzer et al. 2007; Lashuel et al. 2002; Villar-Pique et al. 2016; Vicente Miranda et al. 2017), or the process of aggregation itself (Oueslati, Fournier, and Lashuel 2010; Taschenberger et al. 2012; Reynolds et al. 2017), with the thermodynamic end-product of aggregation - the amyloid fibril - possibly being involved in the cell-to-cell transmissibility of the disease phenotype (Thakur et al. 2017).

The artificial, highly toxic, but not synucleinopathy-related mutant E57K (Winner et al. 2011) is interesting to mention in the context of the 3D structure presented, because E57 is also at the inter-molecular interface (Figure 2). The presence of a positive side chain at this position in E57K would significantly interfere with the formation of the interface and even the amyloid fibril (Winner et al. 2011). Indeed, this mutant was designed in a successful structure-based attempt to interfere with amyloid fibril formation (at least under some conditions) (Winner et al. 2011). Furthermore, both in a lentivirus-rat system as well as in a transgenic mouse model, the E57K mutant formed a significant amount of oligomers and was highly toxic, resulting in a large decay of TH-sensitive neurons in the *Substantia nigra* of rats and a motor phenotype reminiscent of PD in mice (Winner et al. 2011). Thus, the artificial mutant E57K can be regarded as severe “familial PD-like” mutation both from the *in vivo* and from the structure/mechanism-based point of view.

### Comparison with earlier structural data

Full-length α-Syn subunits in a fibril studied by NMR were found to form a Greek key-like motif (Figure 4A) (Tuttle et al. 2016), which is consistent with the here reported structure of α-Syn(1-121). However, the fibrils used for the NMR study were approximately 5 nm wide, which corresponds to the diameter of a single protofilament. The larger diameter of our fibrils, 10 nm, results from the interaction between two protofilaments, which allowed us to elucidate the nature of α-Syn(1-121) protofilament aggregation. Fibrils of 5 to 10 nm in diameter have been previously found in *substantia nigra* samples from the brain of PD patients (Crowther, Daniel, and Goedert 2000). Interestingly, in our structure residue A53, which mutation to threonine is associated with early onset PD, forms part of the interface between two protofilaments, while in the NMR structure of full-length human α-Syn fibrils (Tuttle et al. 2016), residue A53 does not face the inter-molecular interface but points towards the hydrophobic core of individual protofilaments, which may explain the lack of 10 nm fibrils in their sample.

**Figure 4.**
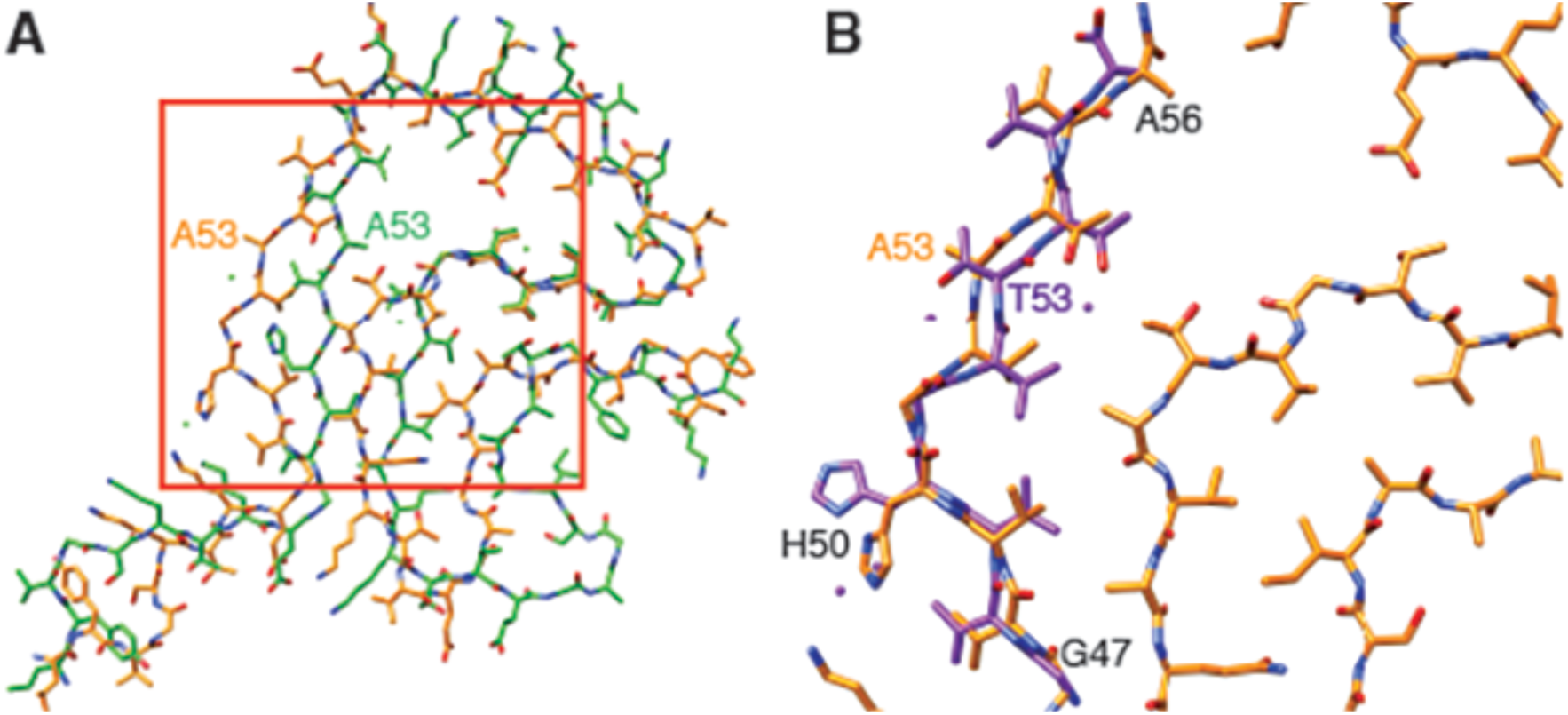
Comparison of α-Syn(1-121) fibrils with previous α-Syn fibril structures. **(A)** Overlay with the solid-state NMR structure from Tuttle, *et al.*(Tuttle et al. 2016) (green). Our α-Syn structure is orange in both overlays. **(B)** Overlay with the preNAC segment obtained by micro-ED by Rodriguez, *et al.* (Rodriguez et al. 2015) (purple). The red square in (A) indicates the area of our structure shown in (B). Residue 53 is mutated (i.e., A53T) in the micro-ED structure.

Our structure includes a serine residue at position 87 (Figure 1 – figure supplement 3E), which is one of the several phosphorylation sites in α-Syn, in addition to Y125, S129, Y133 and Y135 (Oueslati et al. 2012; Paleologou et al. 2010). S87 is the only phosphorylation site located within the NAC region. The previous solid-state NMR structure of α-Syn placed the side chain of this residue towards the inside of the protofilament core, leading to the assumption that phosphorylation of S87 might be the only modification occurring at a region not accessible in the fibrillar state. However, in our cryo-EM structure, S87 faces the outside of the fibril and hence remains accessible for disease-associated modification in α-Syn fibrils.

We also observed the arrangement of G47 and A78 described by Tuttle *et al.* (Tuttle et al. 2016), which was proposed to favor the interaction between residues E46 and K80 and allow them to form a stable salt bridge between two consecutive α-Syn monomers (Figure 1 – figure supplement 3G). The conservation of the geometry adopted by these residues confirms their role in facilitating backbone-backbone interactions. In addition, our structure also confirms that residues A69 and G93 (and likely G68) help to stabilize the distal loop in a protofilament (Figure 1 – figure supplement 3F).

A microED structure obtained from crystals produced from a 10-residue peptide simulating the core of α-Syn fibrils (PreNAC, from 47 to 56; Figure 4B) and including a threonine at position 53, also proposed that residue 53 forms the hydrophobic core within a protofilament (Rodriguez et al. 2015). In addition, the microED model suggested that the interaction between adjacent protofilaments would occur through residues 68 to 78 (referred to as NACore) (Rodriguez et al. 2015). However, their short peptides did not include most residues responsible for the Greek key-like topology that we observed. Instead, our cryo-EM structure reveals that the PreNAC is responsible for the interaction between protofilaments, and places the NACore at the very center (*i.e.*, the core) of a single protofilament.

### Possible mechanism of fibril growth

Based on the above discussion, the fibril structure presented here is likely to be the transmissible species, since an α-Syn polymorphism with a similar Greek key-like motif has the ability to initiate a disease-like behavior (Tuttle et al. 2016). Our 3D structure thereby reveals some detailed insight into the mechanism of fibril replication. Because two different stacking modes are present (*i.e.*, the half-stack at the intermolecular interface and the stacking of β-strand β6), the two ends of the fibrils are distinct, suggesting an end-dependent growth of the fibrils, as documented and also suggested for other amyloids (Luhrs et al. 2005). One end of the fibrils includes a hydrophobic cleft formed between β-strands β2/ β3 on one side and β7 on the other side (residues V49, V52, A88, I89), providing a hydrophobic entry point for the next incoming molecule, with the matching segment (residues V74-V82) including 5 hydrophobic residues (Figure 5). This indicates that the initial binding event of fibril elongation might be a hydrophobic interaction involving residues V74-V82. This peptide segment is the central part of the NAC region and strong experimental evidence suggests that it is critical for fibril formation (Giasson et al. 2001). In addition, it has been shown that β-synuclein, which lacks residues V74 to V82, is mostly or entirely incapable of forming fibrils (Giasson et al. 2001).

**Figure 5.**
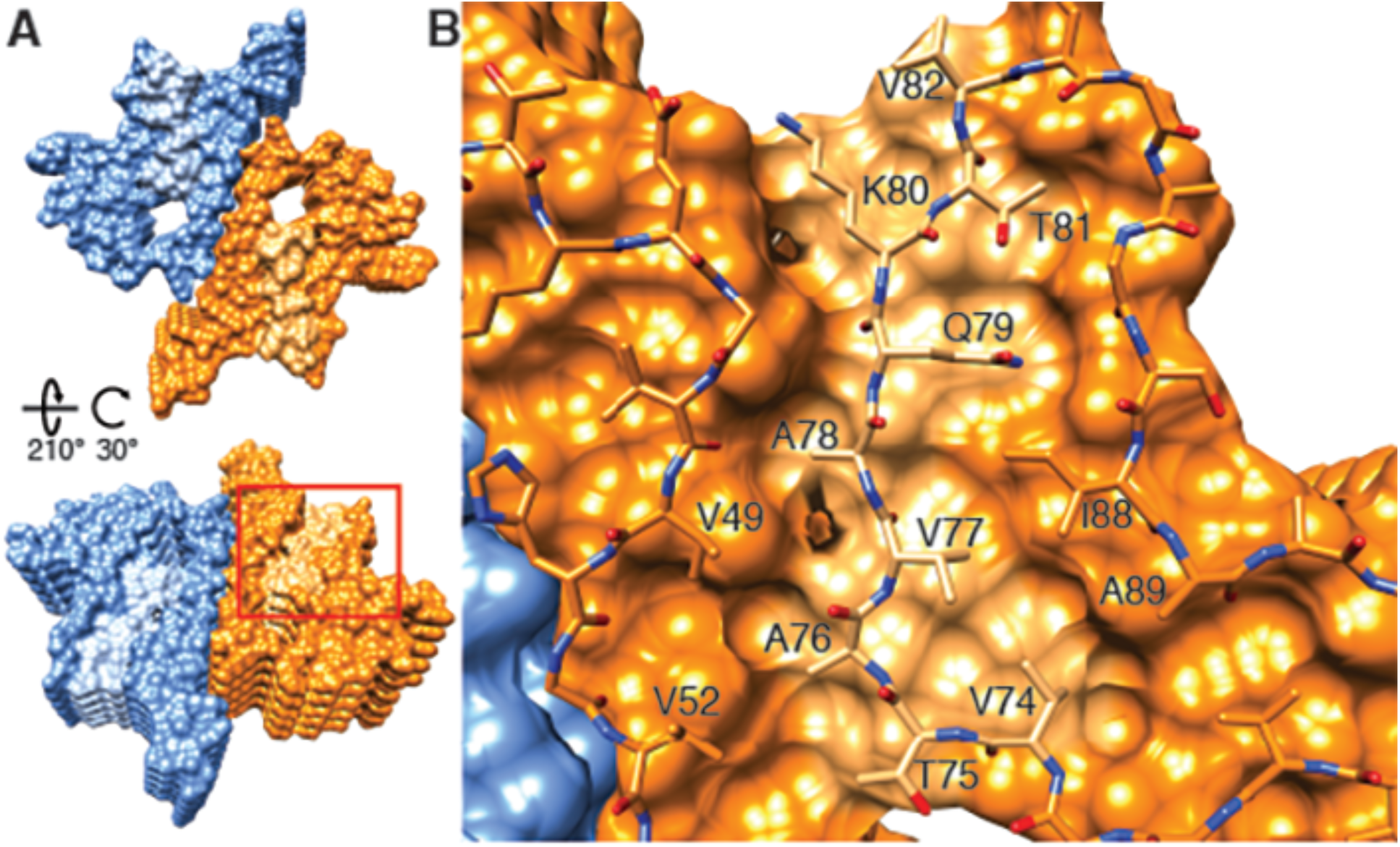
Hydrophobic cleft at the growing end of α-Syn(1-121) fibrils. **(A)** Views of opposite ends of α-Syn fibrils with the two protofilaments colored orange and blue. Regions corresponding to the location of the hydrophobic cleft are shown in a lighter shade. **(B)** Residues forming the hydrophobic cleft, including V49, V52, I88, A89 provide an entry point for residues V74-V82 of an incoming α-Syn molecule (atoms shown). Area shown in panel (B) is marked in panel (A) with a square.

It is intriguing to speculate that a small molecule binding into this hydrophobic cleft would be a potent fibril growth inhibitor or tracer, with the potential to be applied in PD and other synucleinopathies. Finally, the inter-molecular stacking may also play a role in fibril elongation, since the zipper interaction is of hydrophobic nature. Furthermore, it is likely that fibril growth alternates between the two protofilament structures at the level of monomer addition. Failure thereof would possibly result in the growth of a single protofilament with little stability, yielding a dynamic on- and off-binding of monomers and larger oligomers, which has been observed for other amyloid fibril systems (Carulla et al. 2005).

In conclusion, we present the structure of recombinant α-Syn(1-121) fibrils determined at a resolution of 3.4 Å by cryo-EM. Our structure encompasses nearly the complete protein, including the NAC region (residues 61 to 95) of α-Syn. We determined that various residues associated with familial forms of PD and other synucleinopathies are located in the interacting region between two protofilaments, suggesting their involvement in fibril formation and stabilization. The cryo-EM structure presented here reveals how two protofilaments interact to form a fibril, and how the NAC region contributes to protofilament formation and stability. Our structure also presents novel insights into how several PD-relevant mutations of α-Syn would compromise the structure of this fibril, suggesting that a different structure of α-Syn than this fibril strain might be the causative agent in familial PD. Our findings on the mechanism of fibril elongation and protofilament interaction open the door to the informed design of molecules for diagnostics or treatment of synucleinopathies.

## Materials and Methods

### Recombinant proteins

Recombinant full-length α-Syn was expressed from the pRT21 expression vector in BL21(DE3) competent *Escherichia coli* (*E. coli*). For N-terminal acetylation of α-Syn, cells were pre-transfected by pNatB vector coding for the N-terminal acetylase complex (plasmid kindly provided by Daniel Mulvihill, School of Biosciences, University of Kent, Canterbury, UK) (Johnson et al. 2010). C-terminally truncated forms of α-Syn(1-119), α-Syn(1-121), and α-Syn(1-122) were expressed in BL21-DE3-pLysS competent *E. coli* (plasmids courtesy of Prothena Biosciences, South San Francisco, CA, USA). Purification of α-Syn strains was performed by periplasmic lysis, ion exchange chromatography, ammonium sulfate precipitation, and gel filtration chromatography as previously described (Huang et al. 2005; Luk et al. 2009). Polo like kinase 2 (PLK2) was expressed in BL21-DE3-pLysS competent *E. coli*, isolated via its His-tag and immediately used to phosphorylate purified α-Syn. This was followed by standard ion exchange and gel filtration chromatography to separate phosphorylated from non-phosphorylated α-Syn. Endotoxins were removed from all α-Syn strains by Detoxi-Gel Endotoxin Removing Gel (Thermo Scientific) usually in one run or until endotoxin levels were below detection level. The sequence of the expressed α-Syn strains was verified by tryptic digestion followed by MALDI mass spectrometry (MS) or HPLC/ESI tandem MS for total mass was performed. Purity and monodispersity was determined by Coomassie blue or Silver staining of the SDS PAGE gel and analytical ultracentrifugation and the concentration was determined by the bicinchoninic acid (BCA) assay (Thermo Scientific) with bovine serum albumin as a standard. Dialyzed and lyophilized α-Syn(1-121) was prepared by dialyzing the purified protein in a 2 kD Slide-A-Lyzer unit (Thermo Scientific, for max. 3 ml) against HPLC-water (VWR). 500 μg protein aliquots were pipetted into 1.5ml tubes, frozen on dry ice, and lyophilized for 2h using an Eppendorf concentrator (Eppendorf). Lyophilized samples were stored at −80°C until use.

### Fibrillization

Fibrils were prepared by dissolving dialyzed and lyophilized, recombinant α-Syn protein at 5 mg/mL in incubation buffer (DPBS, Gibco; 2.66mM KCL, 1.47mM KH_2_PO4, 137.93mM NaCl, 8.06mM Na_2_HPO_4_-7H_2_O pH 7.0 – 7.3). Reactions of 200 μL per tube were incubated at 37°C with constant agitation (1,000 rpm) in an orbital mixer (Eppendorf). Reactions were stopped after 5 days, sonicated (5 min in a Branson 2510 water bath), aliquoted, and stored at −80°C until use. The presence of amyloid fibrils was confirmed by thioflavin T fluorimetry and high molecular weight assemblies were visualized by gel electrophoresis.

### Electron microscopy

Cryo-EM grids were prepared using a Vitrobot Mark IV (ThermoFisher Scientific) with 95% humidity at 4 °C. Amyloid fibrils (3 μL aliquots) were applied onto glow-discharged, 300 mesh, copper Quantifoil grids. After blotting, grids were plunge frozen in liquid ethane cooled by liquid nitrogen. Samples were imaged on a Titan Krios (ThermoFisher Scientific) transmission electron microscope, operated at 300 kV and equipped with an energy filter (GIF, 20eV energy loss window; Gatan Inc.). Images were acquired on a K2 detector (Gatan Inc.) in dose fractionation mode (50 frames) using the Serial EM software (Mastronarde 2005) at a magnification of 165,000 × (physical pixel size 0.831 Å) and a total dose of ∼69 electrons per square angstrom (e^-^/Å^2^) for each micrograph. Micrographs were drift-corrected and dose-weighted using MotionCor2 (Zheng et al. 2017) through the Focus interface (Biyani et al. 2017). Additional data collection parameters are detailed in Table 1.

**Table 1.**
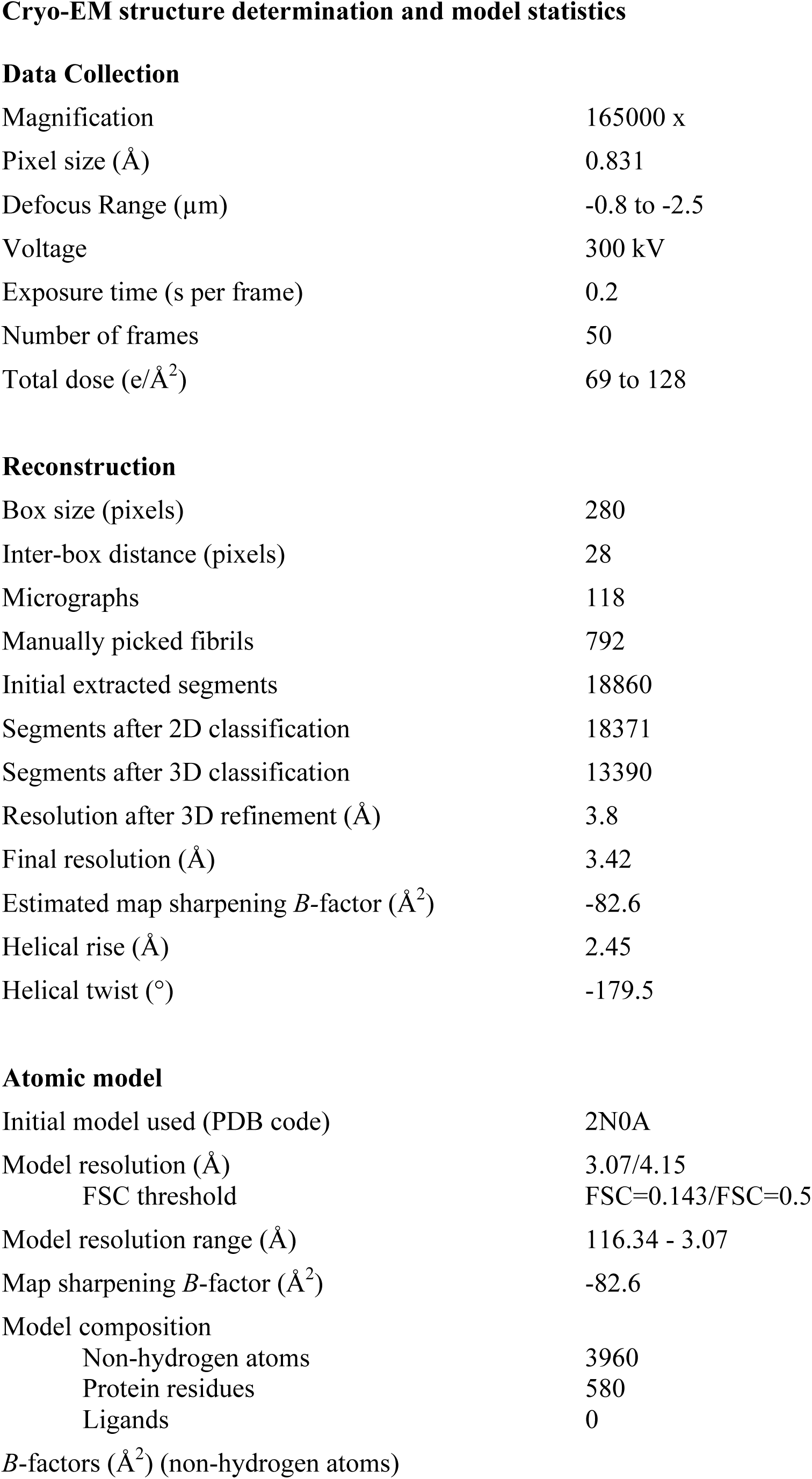
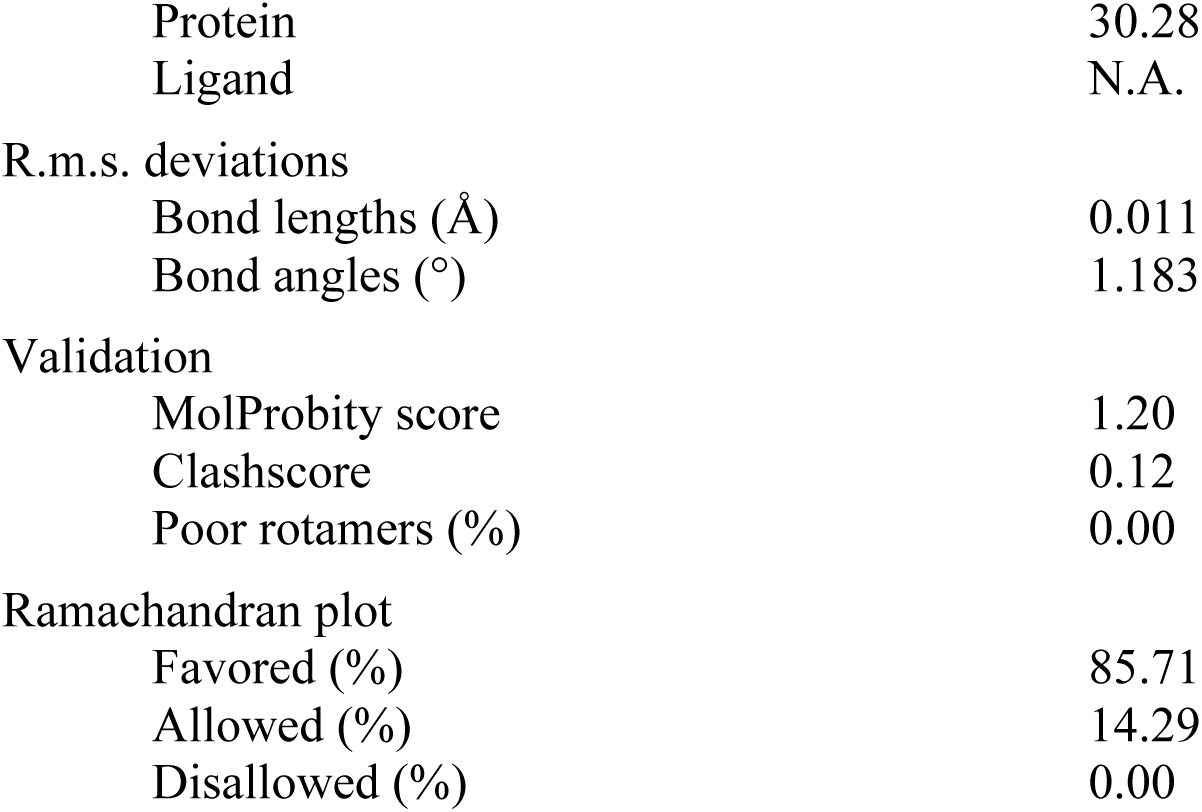
Cryo-EM structure determination and model statistics.

**Data Collection**
|  |  |
| --- | --- |
| Magnification | 165000 x |
| Pixel size (Å) | 0.831 |
| Defocus Range (μm) | -0.8 to -2.5 |
| Voltage | 300 kV |
| Exposure time (s per frame) | 0.2 |
| Number of frames | 50 |
| Total dose (e/Å<sup>2</sup>) | 69 to 128 |
| **Reconstruction** |  |
| Box size (pixels) | 280 |
| Inter-box distance (pixels) | 28 |
| Micrographs | 118 |
| Manually picked fibrils | 792 |
| Initial extracted segments | 18860 |
| Segments after 2D classification | 18371 |
| Segments after 3D classification | 13390 |
| Resolution after 3D refinement (Å) | 3.8 |
| Final resolution (Å) | 3.42 |
| Estimated map sharpening *B*-factor (Å<sup>2</sup>) | -82.6 |
| Helical rise (Å) | 2.45 |
| Helical twist (°) | -179.5 |
| **Atomic model** |  |
| Initial model used (PDB code) | 2N0A |
| Model resolution (Å) | 3.07/4.15 |
| FSC threshold | FSC=0.143/FSC=0.5 |
| Model resolution range (Å) | 116.34 - 3.07 |
| Map sharpening *B*-factor (Å<sup>2</sup>) | -82.6 |
| Model composition |  |
| Non-hydrogen atoms | 3960 |
| Protein residues | 580 |
| Ligands | 0 |
| *B*-factors (Å<sup>2</sup>) (non-hydrogen atoms) |  |
| Protein | 30.28 |
| Ligand | N.A. |
| R.m.s. deviations |  |
| Bond lengths (Å) | 0.011 |
| Bond angles (°) | 1.183 |
| Validation |  |
| MolProbity score | 1.20 |
| Clashscore | 0.12 |
| Poor rotamers (%) | 0.00 |
| Ramachandran plot |  |
| Favored (%) | 85.71 |
| Allowed (%) | 14.29 |
| Disallowed (%) | 0.00 |

### Image processing

Helical reconstruction was carried out in the RELION 2.1 software (Scheres 2012), using methods described by He and Scheres (He and Scheres 2017). Filaments were manually selected using the helix picker in RELION 2.1. Filament segments were extracted using a box size of 280 pixels (233 Å) and an inter-box distance of 28 pixels. A total of 18,860 segments were extracted from 792 fibrils manually picked from 118 micrographs (Table 1). 2D classification was carried out with a regularization value of T=10, and 2D class averages with a clear separation of β-strands were selected for further data processing. Segments assigned to the best 2D classes were used for 3D classification using a regularization value of T=8 and with optimization of the helical twist and rise. For both 3D classification and refinement, a *helical_z_percentage* parameter of 10% was used, which defines the size of the central part of the intermediate asymmetrical reconstruction that is used to apply real-space helical symmetry (He and Scheres 2017). An initial reconstruction was calculated using a cylinder generated via the helix toolbox in RELION 2.1 as initial model. This reconstruction was low-pass filtered to 60 Å and employed as the initial model for a 3D classification with a single class (K=1) and T=20, an approach that allowed the successful reconstruction of amyloid filaments (Fitzpatrick et al. 2017).

Refinement was carried out by the auto-refine procedure with optimization of helical twist and rise. This resulted in a structure with overall resolution of 3.8 Å. Post-processing with a soft-edge mask and an estimated map sharpening *B*-factor of −82.6 Å gave a map with a resolution of 3.4 Å (by the FSC 0.143 criterion). An estimation of local resolution was obtained using RELION 2.1 and a local-resolution-filtered map was calculated for model building and refinement.

### Model building and refinement

A model of the α-Syn(1-121) fibril was built into the Relion local resolution-filtered map using COOT (Emsley and Cowtan 2004), with the PDB ID 2N0A as an initial model for the early interpretation of the map. The structure helped to determine the directionality of the protein chain and facilitated the assignment of densities in the map to specific residues. However, due to the large differences between the NMR structure and our EM map, major rebuilding was necessary. The high quality of the EM map allowed us to unambiguously build residues 38-95. A comparison was also carried out between our structure and X-ray structures of α-Syn fragments 69-77 (PDB ID 4RIK), 68-78 (PDB ID 4RIL) and 47-56 (PDB ID 4ZNN; with the mutation A53T).

The structure (10 monomers, 5 on each protofilament) was refined against the RELION local resolution-filtered map with PHENIX real space refine (Afonine et al. 2013). Rotamer, Ramachandran restraints, and “NCS” constraints were imposed, and two *B*-factors per residue were used during refinement. For validation, we randomized the coordinates (with a mean shift of 0.3 Å) and refined (using the same settings) against one of the refinement half-maps (half-map 1). We then calculated the FSC between that model (after refinement against half-map 1) and half-map 1, as well as the FSC between the same model and half-map 2 (against which it was not refined). The lack of large discrepancies between both FSC curves indicates no overfitting took place.

## Acknowledgments

We thank Liz Spycher, Jana Ebner, Alexandra Kronenberger, Daniel Schlatter, Daniela Huegin, Ralph Thoma, Christian Miscenic, Martin Siegrist, Sylwia Huber, Arne Rufer, Eric Kusznir, Peter Jakob, Tom Dunkley, Joerg Hoernschmeyer, and Johannes Erny at Roche for their technical support to clone, express, purify and characterize the different recombinant forms of α-Syn, Kenneth N. Goldie, Lubomir Kovacik and Ariane Fecteau-Lefebvre for support in cryo-EM, and Shirley A. Müller for support in manuscript preparation, and Sjors Scheres for help in image processing. Calculations were performed using the high-performance computing (HPC) infrastructure administered by the scientific computing center at University of Basel (sciCORE; http://scicore.unibas.ch). This work was in part supported by the Synapsis Foundation Switzerland, and the Swiss National Science Foundation (grants CRSII3_154461 and CRSII5_177195). The authors declare no competing financial interests.

---

**Video 1**

**Cryo-EM structure of alpha-synuclein fibril.** Details of the cryo-EM reconstruction of an alpha-synuclein fibril at 3.4 Å resolution, illustrating the interaction between two protofilaments, the 4.9 Å spacing between β-strands of a single protofilament and the Greek-key-like motif in the protofilament core.

**Figure 1 – figure supplement 1.**
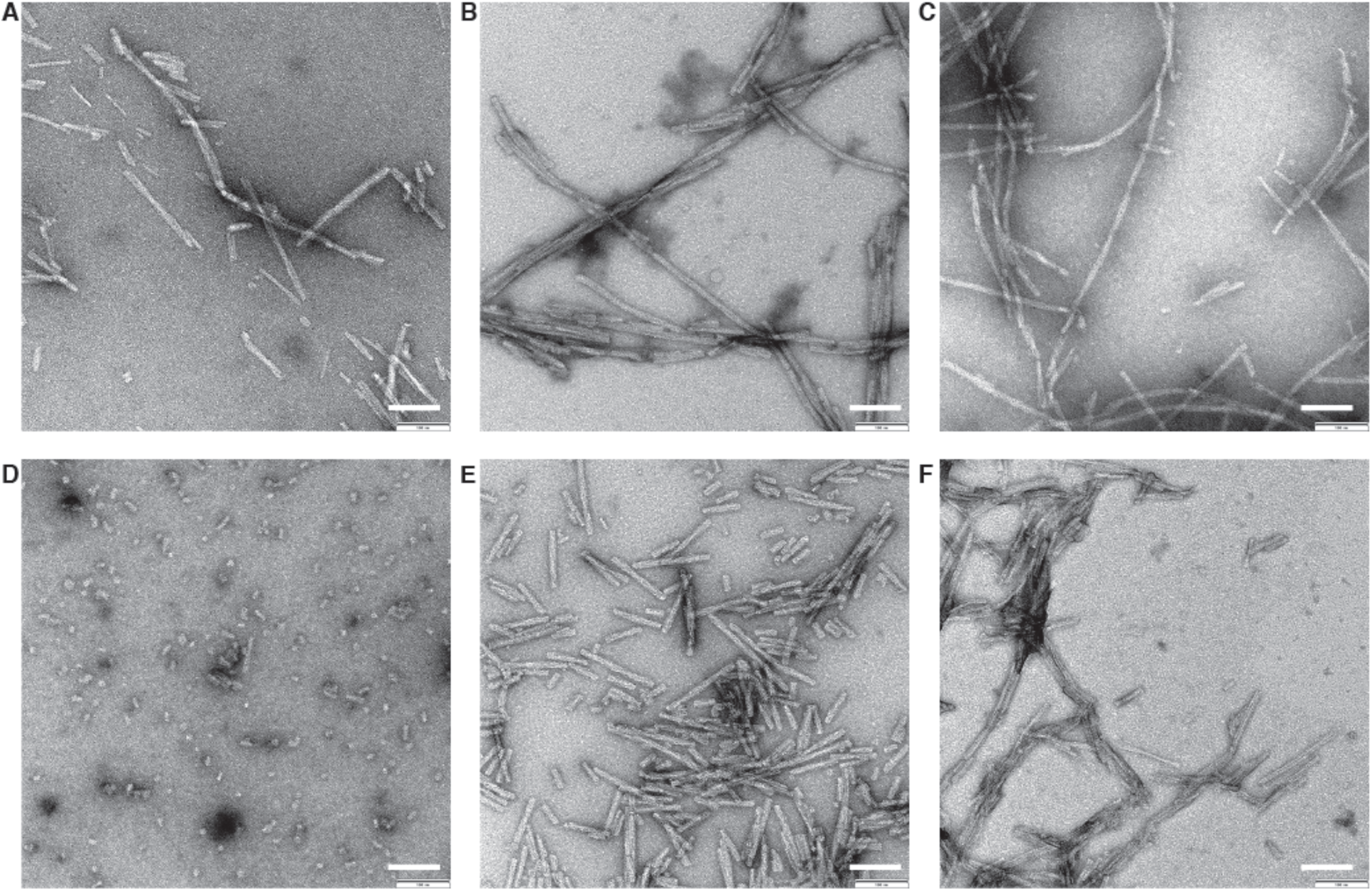
Negative stain TEM images of α-Syn strains. **(A)** Wildtype. **(B)** Ser129 Phosphorylated. **(C)** N-terminally acetylated. **(D)** C-terminally truncated (α-Syn(1-119)). **(E)** C-terminally truncated (α-Syn(1-121)). **(F)** C-terminally truncated (α-Syn(1-122)). 0.5 mg/ml fibril preparations were stained with 2% uranyl acetate. Scale bars: 100 nm.

**Figure 1 – figure supplement 2.**
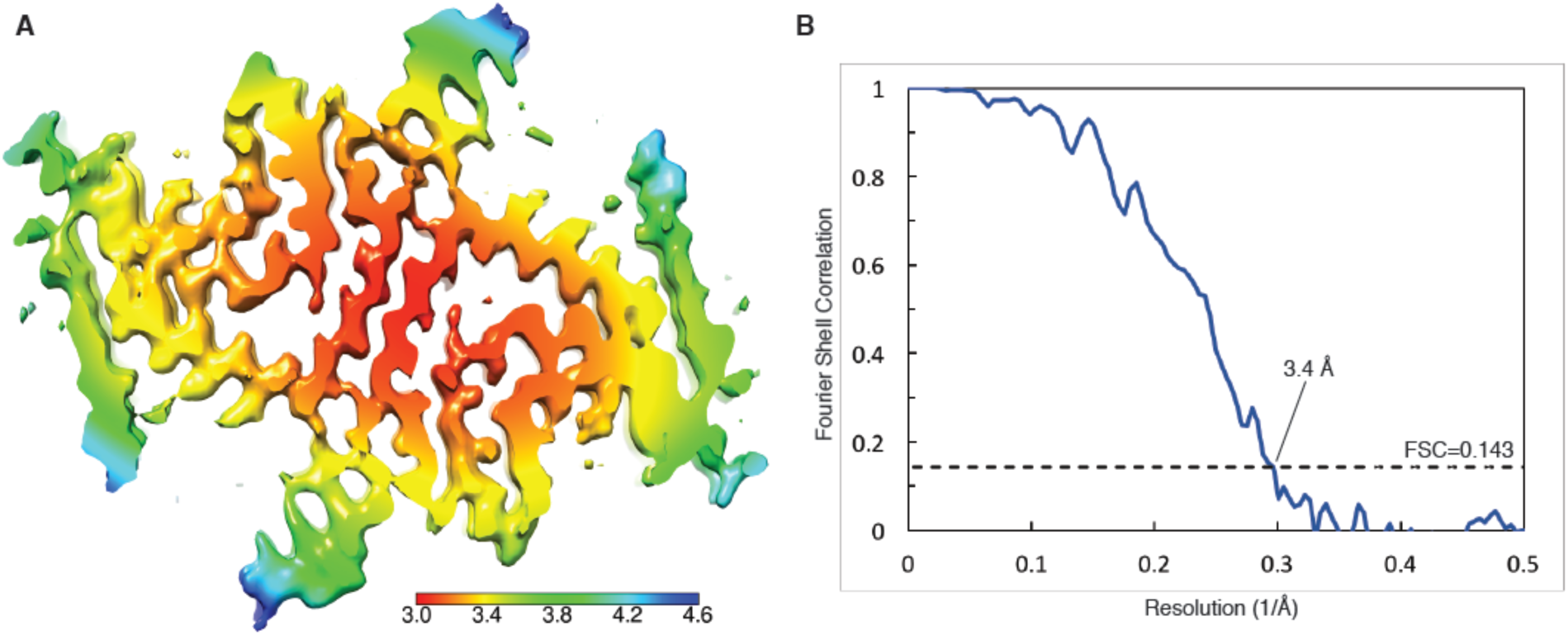
Local resolution estimation and FSC curves. **(A)** Cryo-EM map with local resolution estimation; the color scale indicates resolution ranging from 3.0 Å to 4.6 Å. **(B)** Fourier shell correlation curve between two independently-refined half-maps, indicating an overall resolution of 3.4 Å.

**Figure 1 – figure supplement 3.**
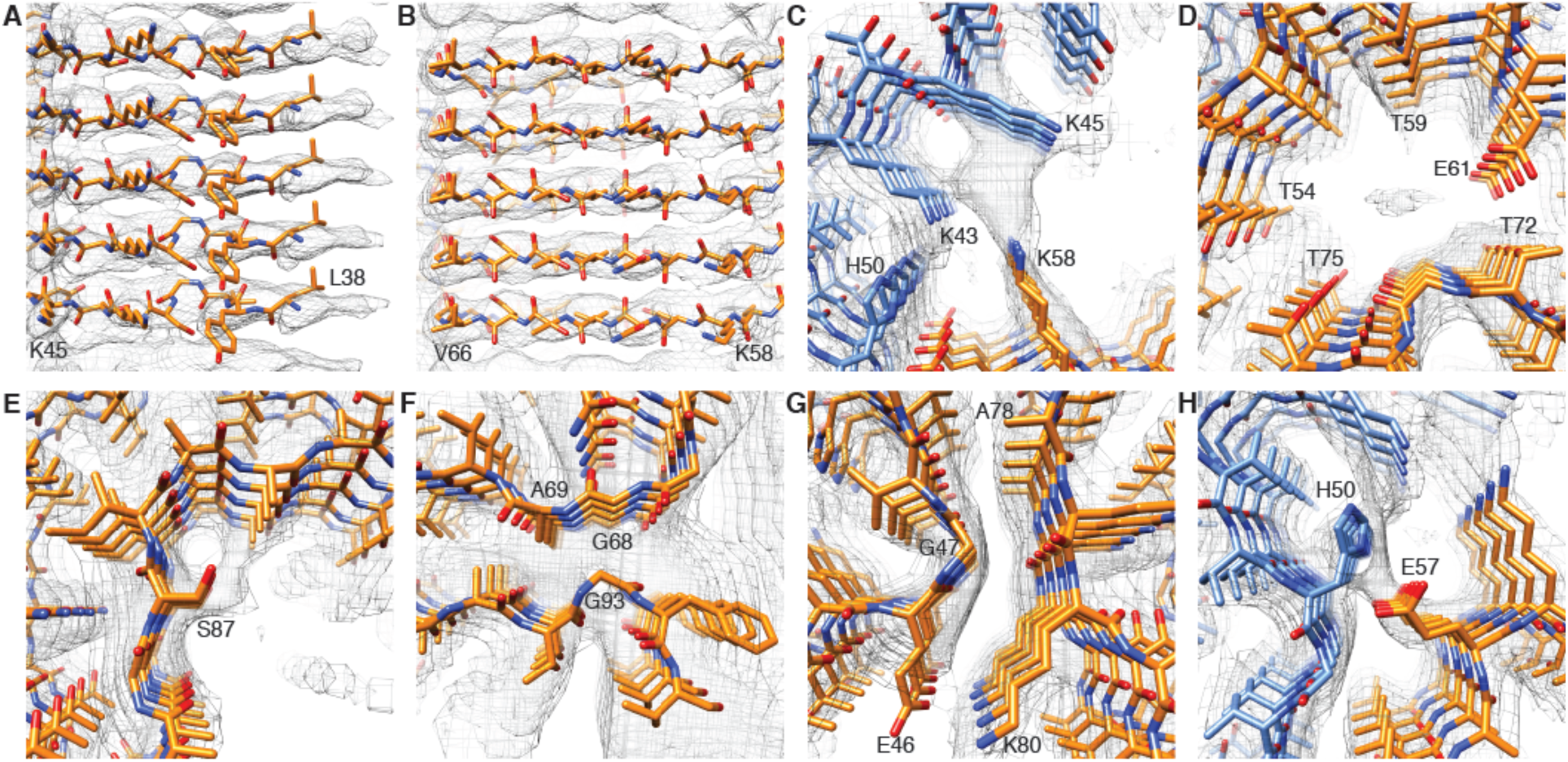
Details of atomic model and density. **(A) (B)** Clearly resolved separation of individual β-strands along the fibril. **(C)** Extra density at the interface of adjacent protofilaments between the positively charged lysines K43 and K45 and lysine K58. **(D)** Hydrophilic region surrounding a tunnel filled with an electron density**. (E)** Phosphorylation site S87 showing the location of this residue towards the outside of the fibril. **(F)** Distal loop of a protofilament indicating residues G68, A69 and G93, which may contribute to loop stability. **(G)** Arrangement of G47 and A78, which may contribute to the interaction between E46 and K80. **(H)** Interaction between H50 and E57 in the interface region of two protofilaments, which may contribute to the stability of protofilament interaction.

**Figure 2 – figure supplement 1.**
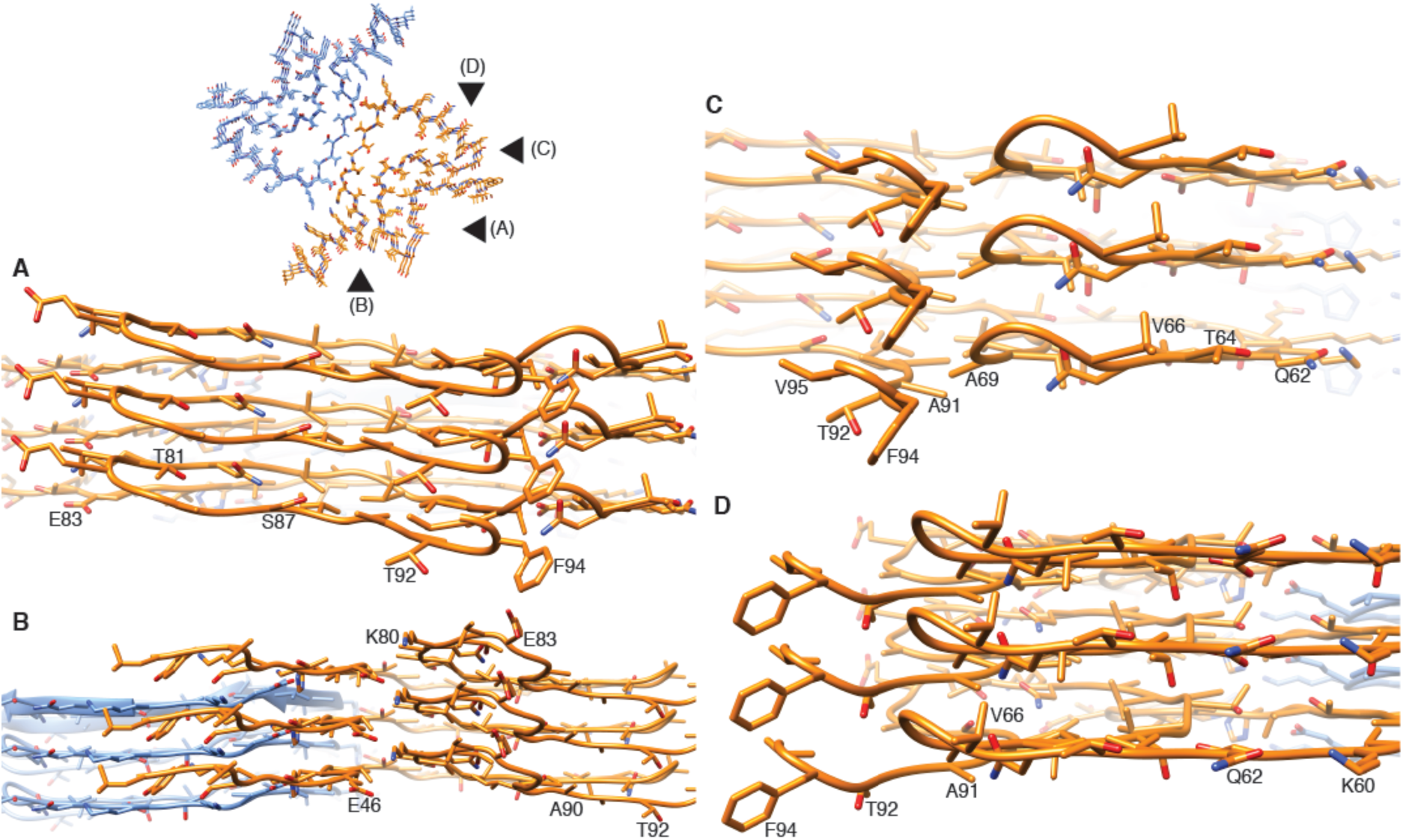
Stacking of β-strands. **(A)-(D)** Close-up side views of the α-Syn fibril illustrating homo- and hetero-steric zippers present in the structure. Pointers in the cross-section panel (top left) indicate the points of view for panels (A) to (D).

